# A novel behavioral paradigm using mice to study predictive postural control

**DOI:** 10.1101/2024.07.01.601478

**Authors:** Yurika Doi, Meiko Asaka, Richard T. Born, Dai Yanagihara, Naoshige Uchida

## Abstract

Postural control circuitry performs the essential function of maintaining balance and body position in response to perturbations that are either self-generated (e.g. reaching to pick up an object) or externally delivered (e.g. being pushed by another person). Human studies have shown that anticipation of predictable postural disturbances can modulate such responses. This indicates that postural control could involve higher-level neural structures associated with predictive functions, rather than being purely reactive. However, the underlying neural circuitry remains largely unknown. To enable studies of predictive postural control circuits, we developed a novel experimental paradigm for *mice*. In this paradigm, modeled after human studies, a dynamic platform generated reproducible translational perturbations. While mice stood bipedally atop a perch to receive water rewards, they experienced backward translations that were either unpredictable or preceded by an auditory cue. To validate the paradigm, we investigated the effect of the auditory cue on postural responses to perturbations across multiple days in three mice. These preliminary results serve to validate a new postural control experimental paradigm, opening the door to the types of neural recordings and circuit manipulations that are currently possible in mice.

**Significance Statement:** The ability to anticipate disturbances and adjust one’s posture accordingly—known as “predictive postural control”—is crucial for preventing falls and for advancing robotics. While human balance is often assessed via floor perturbations, rodent studies typically use rotarod tests. Here, we developed a postural perturbation task for freely moving mice, modeled after those used in human studies. Using a dynamic platform, we delivered reproducible perturbations with or without preceding auditory cues and quantified how the auditory cue affects postural responses to perturbations. Our work provides validation of a new postural control experimental paradigm, which opens the door to the types of neural population recordings and circuit manipulation that are currently possible in mice.

## Introduction

The ability to leverage prior experience is essential for navigating and interacting with complex, dynamic environments. In motor control, prediction enables feedforward control and helps compensate for sensorimotor feedback delays and sensory noise (Dakin and Bolton, 2018; Wolpert and Flanagan, 2001). Prediction also plays an essential role in postural control – the process of maintaining upright posture and balance – which is vital for daily life. Although traditionally considered a reflexive, sensory-driven system, accumulating evidence shows that *prediction* plays an important role in postural control (Bastian, 2006; Dakin and Bolton, 2018; Jacobs and Horak, 2007). However, neural mechanisms underlying this process remain poorly understood.

To illustrate the role of prediction in postural control, imagine riding a train on your daily commute. As you approach a familiar station, the scenery cues you to anticipate deceleration, prompting you to prepare consciously or subconsciously before the train slows. Your postural responses in this scenario would differ from those triggered by an unexpected deceleration. This type of anticipatory adjustment to a predictable *external* perturbation is referred to as “predictive postural control” (see Table 1 for the categorization of postural adjustments, based on the timing and whether the postural disturbance is external or self-generated).

**Table 1:**
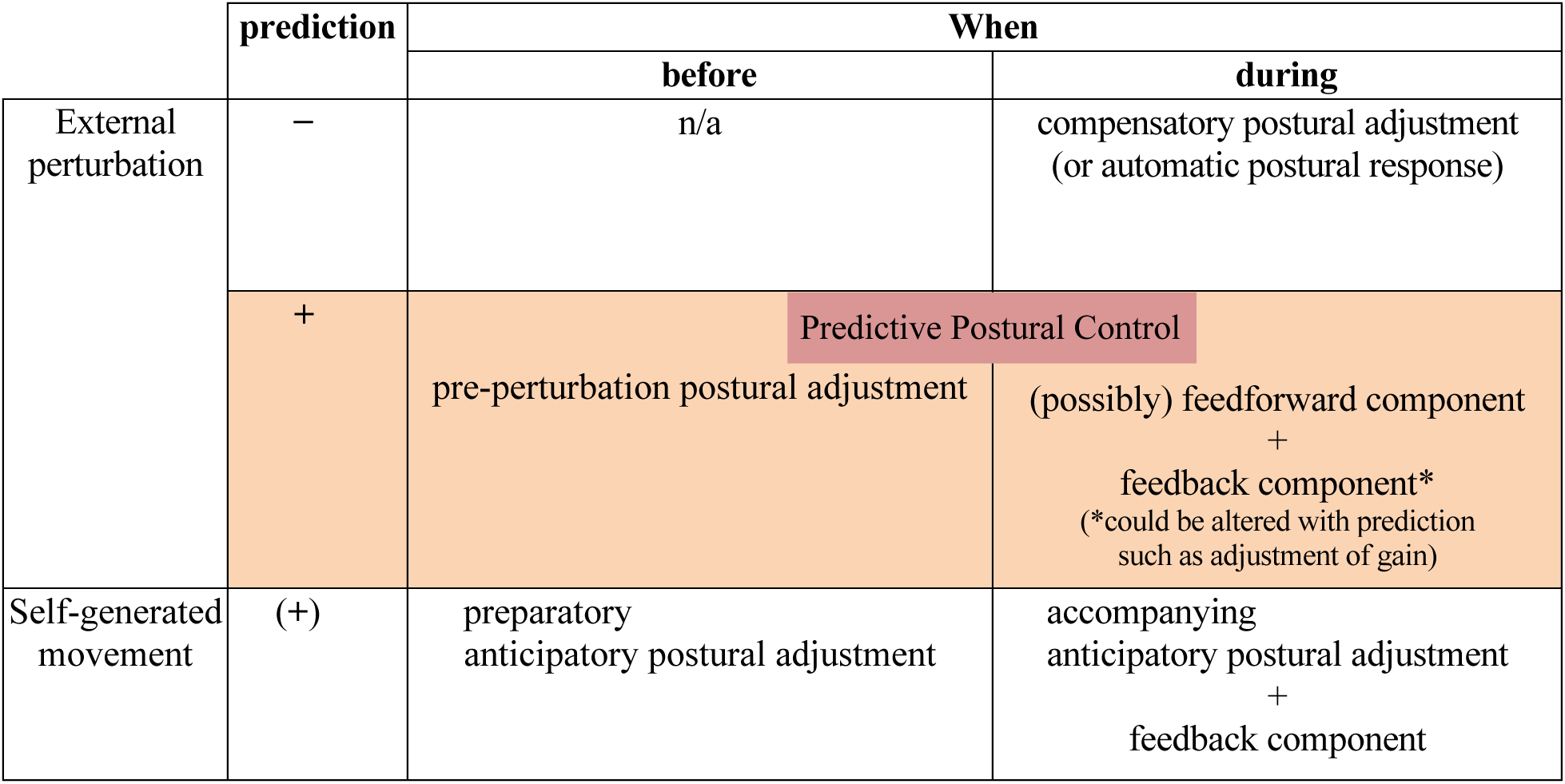
Different Types of Postural Responses (adjustments)

Various behavioral paradigms have been employed to study postural control in humans (Campbell et al., 2009; Coelho et al., 2017; Fujio et al., 2016; Horak et al., 1989; Horak and Diener, 1994; Jacobs et al., 2008; Kolb et al., 2004, 2002; McChesney et al., 1996; Silva et al., 2015). To study *predictive* postural control, several different behavioral paradigms have been used: 1) repetitive patterns of perturbations that rendered them predictable over time (Horak et al., 1989; Horak and Diener, 1994), 2) explicit visual/auditory cues that preceded perturbations and contained some information about the perturbation (e.g. timing and/or direction of the perturbation) (Coelho et al., 2017; Fujio et al., 2016; Jacobs et al., 2008; McChesney et al., 1996; Silva et al., 2015), or 3) classical conditioning, where time-locked coupling of a tone (conditioned stimulus, CS) with a postural perturbation (unconditioned stimulus, US) was repetitively presented (Campbell et al., 2009; Kolb et al., 2004, 2002). Some brain regions such as the cerebellum and the cerebral cortex have been indicated for predictive postural control based on neurological patient studies (Horak and Diener, 1994; Kolb et al., 2004) and electroencephalography (EEG) studies(Jacobs et al., 2008; Mochizuki et al., 2008). However, the precise neural mechanisms remain to be elucidated.

Historically, studies on various non-human animals – including mollusks, lampreys, zebrafish, rodents, and cats – have provided foundational insights into the neural mechanisms of postural control conserved across species (Bagnall and McLean, 2014; Deliagina and Orlovsky, 2002; Macpherson et al., 2007; Schepens and Drew, 2004; Straka and Baker, 2013; Sugioka et al., 2023; Whishaw et al., 1994; Yakovenko and Drew, 2009). However, efforts to understand the neural mechanisms underlying predictive postural control in non-human animals have been limited. Rodent studies have traditionally relied on paradigms using rotarods (Crawley, 2007; Dunham and Miya, 1957) or balance beams (Carter et al., 2001; Luong et al., 2011) to assess general balance control and motor coordination. More recent methods, such as the suspended dowel task (Lee et al., 2015) and the beam destabilization task (Murray et al., 2018), have begun to explore postural control more directly. A paradigm mimicking floor perturbation studies, commonly used in humans, was recently developed for bipedally standing rats, involving floor tilting perturbations preceded by a fixed-interval visual cue (Konosu et al., 2024, 2021). These studies demonstrated reduced postural response amplitude over repeated perturbations, suggesting plasticity in the rat postural control system.

Building on these prior studies, here we sought to establish a novel mouse paradigm to study predictive postural control, modeled after paradigms used in human studies (Horak et al., 1989; Kolb et al., 2002; Welch and Ting, 2014) and recent rat studies (Konosu et al., 2024, 2021). We describe the postural task we developed, quantify postural responses based on kinematics and reward acquisition, and analyze the effects of preceding cues and learning on these responses. This mouse model offers a powerful tool to investigate the neural mechanisms underpinning predictive postural control.

There are multiple ways in which predictive mechanisms can work to affect postural control. One way is the generation of postural adjustments that *precede* an external perturbation (Jacobs et al., 2008; Mochizuki et al., 2008; Welch and Ting, 2014), such as leaning to one side, widening the stance, isometric contractions, etc. Another way is the modulation of the postural response *during* the perturbation. Two possible mechanisms (that are not mutually exclusive) can be considered to produce such modulations. One mechanism is the (feedforward) adjustment of sensorimotor feedback parameters such as the scaling of gains (Lockhart and Ting, 2007; Pruszynski and Scott, 2012). In other words, the control gain of the feedback system can be pre-set before a perturbation occurs. Another possible mechanism is that the feedforward movement command is prepared in advance and gets discharged during the perturbation. In other words, feedforward control can work in parallel with the feedback mechanisms; the idea that is comparable to “accompanying APA (Schepens and Drew, 2004).”

## Methods

### Animals

A total of 3 adult male mice were used. Experiments were performed on C57/BL6 mice supplied by CLEA (Tokyo, Japan), aged 10-26 weeks (27.3 ± 1.2 g baseline pre-restriction weight; two mice [mouse CB5 and CB6] were 13 weeks old at the start of pre-training, 24-26 weeks old during the full task experiments, one mouse [mouse CB10] was 10 weeks old at the start of pre-training, 14-16 weeks old during the full task experiments). Experiments were conducted between 9:00 and 18:00. Animals were individually housed in a room with a constant temperature and a reverse 12-h light and dark cycle. Mice were water-restricted to maintain body weight >85% of their baseline weight to ensure task motivation while maintaining health. Additional water supplementation was provided shortly following experiment sessions with a total of task and post-task water ranging from 0.8 - 1.3mL per day, determined individually per mouse. Body weights ranged from 85 - 108% of baseline pre-restriction weight across mice. All procedures were performed in accordance with the National Institutes of Health Guide for the Care and Use of Laboratory Animals and approved by the Harvard Animal Care and Use Committee. This study was approved by the Ethical Committee for Animal Experiments at the University of Tokyo, and was carried out in accordance with the Guidelines for Research with Experimental Animals of the University of Tokyo. This study was carried out in compliance with the ARRIVE guidelines (Kilkenny et al., 2010; Percie du Sert et al., 2020).

### Predictive postural control task

We developed a postural perturbation task in freely moving mice, modeled after those used in human studies (Horak et al., 1989; Kolb et al., 2002; Welch and Ting, 2014) and a recent rat study (Konosu et al., 2021), in which a dynamic platform was used to give reproducible perturbations.

Water-restricted mice were placed in a clear acrylic box with a perch to stand on to access the lick spout for water reward (Figure 1a). Water restriction enables the use of precise, temporally controlled rewards, which are essential for training mice in nonspontaneous, task-specific behaviors, such as prolonged upright standing. This approach aligns with standard practices in behavioral neuroscience, where operant conditioning is necessary to shape behaviors consistently across trials, ensuring reliability in results and enhancing the relevance of findings across studies. The height of the lick spout was adjusted such that mice could just barely reach it by standing on their two hind legs (bipedal standing) while balancing on the perch. In other words, mice were required to stand on the perch to obtain water reward, which constrained the standing position and body orientation of the mice. Using a bipedal stance provides a simplified experimental model where postural adjustments are more pronounced and easier to measure compared to quadrupedal posture. This would allow us to better isolate and study the fundamental neural circuits involved in postural control. A light-emitting diode (LED) indicated that mice were eligible to start a trial. Mice initiated the trial by licking the spout, which was monitored by a capacitive sensor.

**Figure 1:**
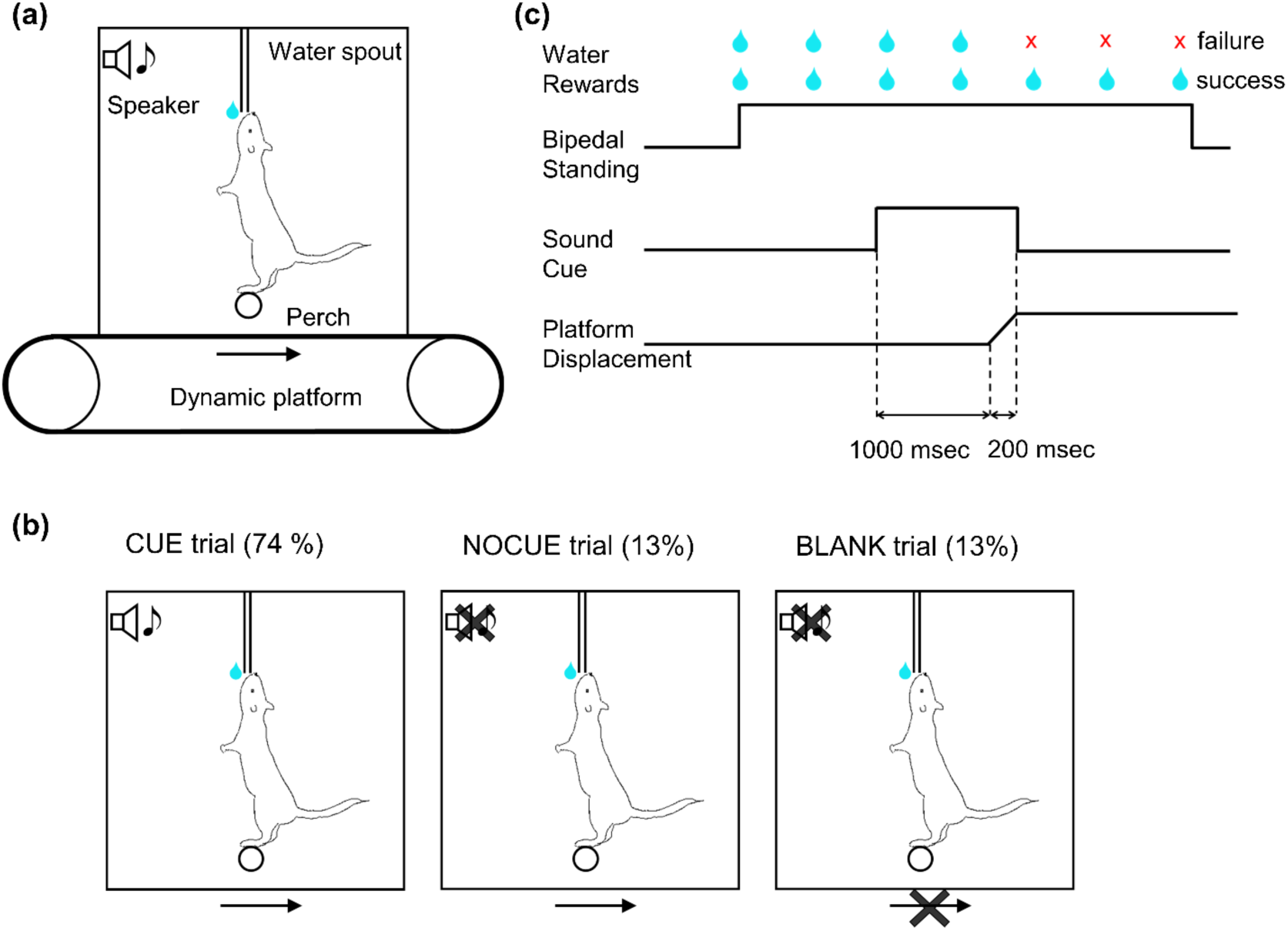
Postural task. (a) Diagram of behavioral apparatus. Mice stood on a round perch to receive water rewards from the lick spout. Both the perch and the water spout were fixed to the behavioral box. The behavioral box was mounted on a dynamic platform that created horizontal movement of the box. (b) Diagram of three different trial types. (1) trials in which a cue preceded platform movement by 1 second. (CUE, 74% of trials); (2) trials in which the platform moved but no cue was given (NOCUE, 13%); and (3) trials with no platform movement (BLANK, 13%) (c) Schematic depicting trial structure in the most common trial type (CUE trials). Mice initiated the trial by licking, which was immediately rewarded with a water droplet (2 µl). Postural perturbations consisted of a backward platform displacement lasting 200 msec. A sound cue was played preceding the perturbation by 1 sec and terminating at the end of the platform movement.

In each trial, a reward water droplet (2 µl) was given immediately after the first lick. A 2 µL water reward was selected to motivate mice effectively without causing rapid satiation, consistent with the typical reward size range in mouse studies (Cohen et al., 2012; Eshel et al., 2016; Histed et al., 2012; Kim et al., 2020). In order to keep the trial active, mice had to continue licking within an interval of 600 milliseconds (msec). Subsequent water droplets were given for each lick that occurred at least 1,100 msec after the previous reward. The height of the lick spout and the requirement of continuous licking, enforced by the maximum 600 msec window with no lick, encouraged the mice not to make large postural changes. The maximum duration of a trial was 7.5 seconds and mice could receive a maximum of 7 water droplets per trial. A trial was terminated in one of two ways: (1) If mice were able to continue licking the spout for the entire 7.5 seconds, the trial was deemed “complete”; (2) If mice failed to lick within any 600-msec interval, the trial was classified as an “aborted” trial. At the end of each trial, the LED was turned off and mice had to wait for an inter-trial duration of 10 - 15 sec (drawn from a uniform distribution) until the next trial could be initiated. The 10-15 second inter-trial interval was intended to provide a brief rest period for the mice, which may minimize fatigue and prevent potential carryover effects. For aborted trials, an additional 20 sec was added to the inter-trial duration as a penalty. In trials with a “perturbation,” mice experienced a backward movement of the entire behavior box, including the perch and the lick spout. A platform displacement occurred after a random delay from the first lick, which was drawn from a truncated exponential distribution (minimum, 2.5 sec; maximum, 6 sec; mean, 1 sec) to minimize the predictability of the timing. Note that if the trial was aborted before the cue or the perturbation, the same trial type was repeated for the next trial.

In order to test whether predictability affects postural responses, we used three different trial types (Figure 1b): (1) trials in which an auditory cue preceded platform movement by 1 second (CUE, 74% of trials); (2) trials in which the platform moved but no cue was given (NOCUE, 13%); and (3) trials with no platform movement (BLANK, 13%). For CUE trials, a 6 kHz tone began 1 sec before the onset of platform movement and terminated at the end of platform movement (Figure 1c). The reason for including BLANK trials is to generate uncertainty about whether the perturbation would happen or not, and to strengthen the relation of perturbation to the cue. All three trial types were randomly interleaved. Each animal participated in only one recording session per day throughout the study period. In each daily session, mice performed 67-119 trials (median = 83 trials). The platform was moved in one of three amplitudes: 7 (small, only used for mouse CB10), 12 (medium), or 18 (large) millimeters. Only one amplitude was used for any single session.

For the sake of brevity, we will refer to perturbation trials (CUE trials and NOCUE trials), which constituted 87% of all trials, simply as “trials”. BLANK trials, which comprised the remaining 13% of trials, will not be mentioned further in the analyses. Note that trial indexing is based on all trial types (including BLANK trials).

#### Behavioral apparatus

Custom hardware and software were used so that the task could run in a semi-automated way with pre-set task parameters using a closed-loop system (Figure 2). Since our paradigm uses an open-source programmable microcontroller that offers a flexible design of the task, it can easily be modified and applied to different postural tasks that ask different questions.

**Figure 2:**
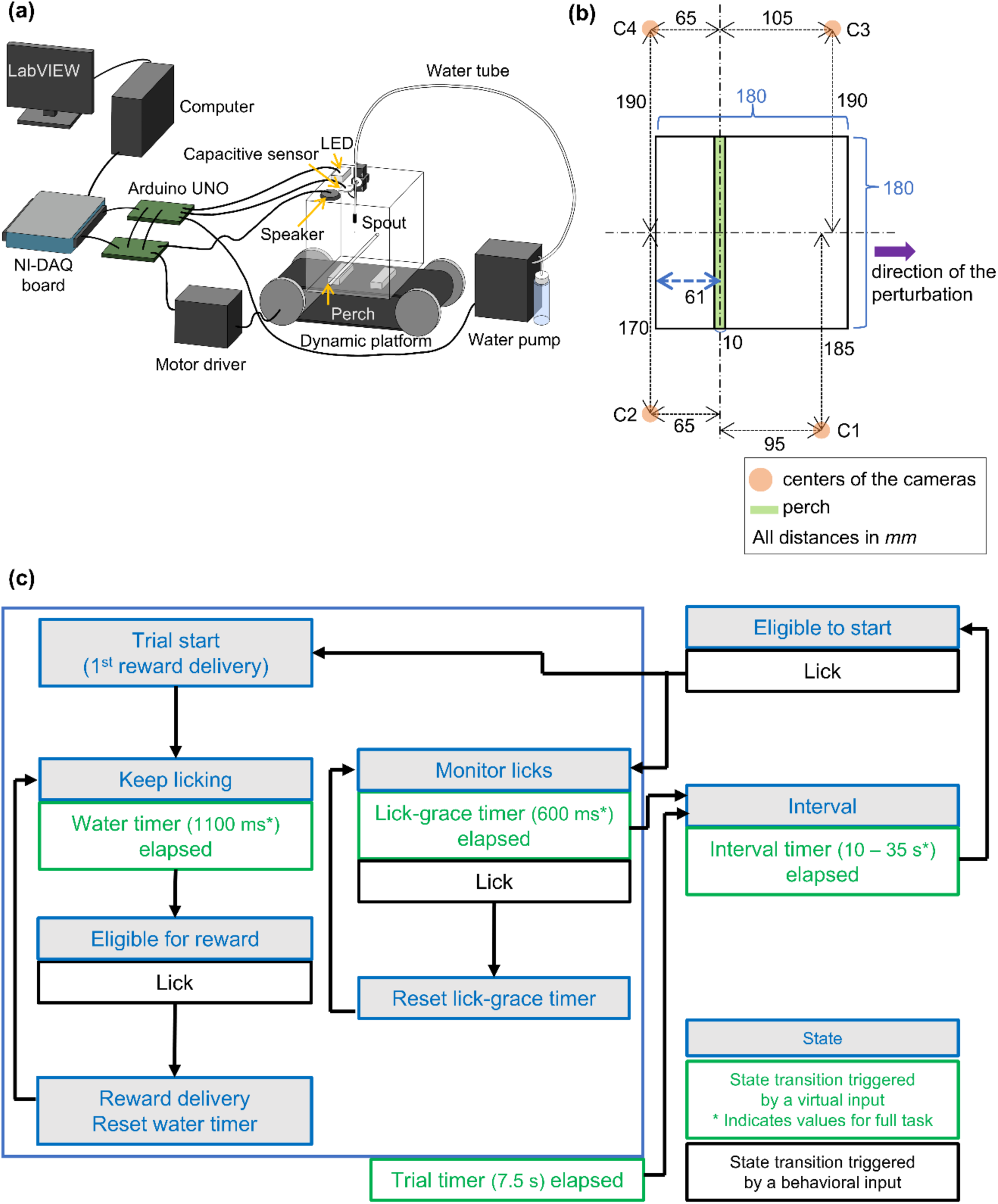
Experiment apparatus and task design. (a) Design of the experimental apparatus. A box made of clear acrylic was mounted on a dynamic platform. A round rod (perch), water spout, LED, and capacitive sensor were fixed to the box. Two microcontrollers (Arduino UNOs) were connected to the electronics to run the task in a closed-loop system. Video cameras and a synchronization device were omitted from the diagram for simplicity. See Table 2 for the parts list. (b) Schematic bird’s-eye view illustrating the 2D spatial arrangement of the cameras relative to the behavioral box. The black rectangle represents the behavioral box. The orange circles (C1, C2, C3, and C4) indicate the center positions of the four cameras, positioned around the box. The green rectangle denotes the perch located in the box. Dashed lines indicate the alignment axes to provide a reference for the relative positioning of the cameras and other elements. The blue double-headed arrow specifies the distance of the perch from the inner wall of the box. The purple arrow indicates the direction of the perturbation. All dimensions are provided in millimeters (mm). (c) Task state machine. States (blue) could transition either by a lick (behavioral input, black) or an elapsed timer (virtual input, green).

**Table 2:** Parts List.

| Part | Name/Description | Link |
| --- | --- | --- |
| Walls/floor | Clear acrylic<br>(thickness: 4mm) | https://www.hazaiya.co.jp/item/19037.html |
| Perch | Clear acrylic rod<br>(diameter: 10mm) | https://www.monotaro.com/g/02476005/ |
| Lick spout | Stainless steel tube (20G) | https://www.nazme.co.jp/product/9-0-serviceinformation/9-5-information/kn-sus-p/ |
| Touch sensor | Capacitive sensor | https://www.adafruit.com/product/1982 |
| Speaker | Full range speaker | https://www.digikey.jp/ja/products/detail/bdnc-holding-limited/BGC-D40-22-4-002/9842990 |
| Audio amplifier | Class D amplifier | https://www.adafruit.com/product/1752 |
| Lighting | LED array (blue) | https://akizukidenshi.com/catalog/g/gI-12344/ |
| Camera | NaturalPoint,<br>Optitrack Prime X 13 | https://optitrack.com/cameras/primex-13/ |
| Synchronization device | NaturalPoint,<br>eSync2 | https://optitrack.com/accessories/sync-networking/esync-2/ |
| Microcontroller | Arduino UNO | https://store.arduino.cc/products/arduino-uno-rev3 |
| Data acquisition system | National Instruments,<br>USB-6002 | https://www.ni.com/en-us/support/model.usb-6002.html |

#### Hardware

A behavioral box was developed that allowed the recording of free movement (Figure 2a, see Table 2 for parts list). Mice were able to move freely in a 180 mm by 180 mm box. An acrylic rod of 10 mm diameter was fixed so that the top of the rod was 24 mm above the floor. Bright blue LED light illuminated the box from above to indicate the trial period. An audio speaker was mounted above the box to produce sound cues. A lick spout was hung from the ceiling of the box, and the height of the spout end was adjusted for individual mice. The lick spout was connected to a capacitive sensor that detected licks. The water pump was calibrated so that 2 μL of water was dispensed from the end of the spout for each water droplet delivery. To record the movement of the mice, four cameras were mounted around the box (Figure 2b). The box was placed on a dynamic platform that can be moved in the horizontal direction in a timed manner. The task was controlled by Arduino UNO microcontrollers (Arduino, Somerville, MA), and task parameters and events were recorded to a computer via a data acquisition board (USB-6002; National Instruments, Austin, TX). Arduino UNOs were programmed to execute the task structure described in the task section and were wired to communicate with electronics such as the capacitive sensor, motor driver, speaker, water pump, and LED.

#### Task implementation

There were two phases in the behavioral paradigm: pre-training and full task. During pre-training, mice learned the trial-interval structure with progressive parameters over sessions. In the full task, mice experienced sound cues and platform motion (hereafter referred to as a “perturbation”), and it was during this phase that we investigated the animals’ postural responses.

##### Pre-training

Before testing mice on the full version of the task in which they experienced perturbations, they were trained on a simpler task. The goal of this pre-training was to train mice to stand bipedally during the trial period and not to stand during the interval period. This trial-interval structure was important for two main reasons: 1) to control the timing of the perturbation relative to the onset of standing so that mice were not standing for a too short or too long period of time before the perturbation, and 2) to motivate mice to perform well by limiting the time when they could obtain water rewards. With the trial-interval structure, if mice failed to perform well in a trial and did not get as many water droplets, they had to wait for the duration of the interval period until the next trial became available to start.

The trial-interval structure was as follows (also see Figure 2c):

A blue LED light turned on to indicate that the mouse was eligible to start a trial. As soon as a lick was registered while the LED was on, the trial started, and the first water droplet was given.

In order to keep the trial active, the mouse had to lick at least once within a 600 msec interval (referred to as “lick-grace timer” in Figure 2c). After each lick, the 600-msec lick-timer was re-started, so that any lick-free period of >600 msec resulted in termination of the trial (see below). However, not every lick was rewarded—a water droplet was only given for licks that occurred at least 1100 msec *after* the previous reward (referred to as “water timer” in Figure 2c). Thus, the maximum number of rewards the mouse could receive on a single trial was 7 drops, but the mouse needs to lick more frequently than this to keep the trial active. Figure 3 shows two trials: the first sequence (panels a-d) shows an example of a “successful” trial; the second (panels e-h) are from a “failed” trial. The terms “successful” and “failed” will be explained in greater detail in the section titled *Trial Outcome*.

**Figure 3:**
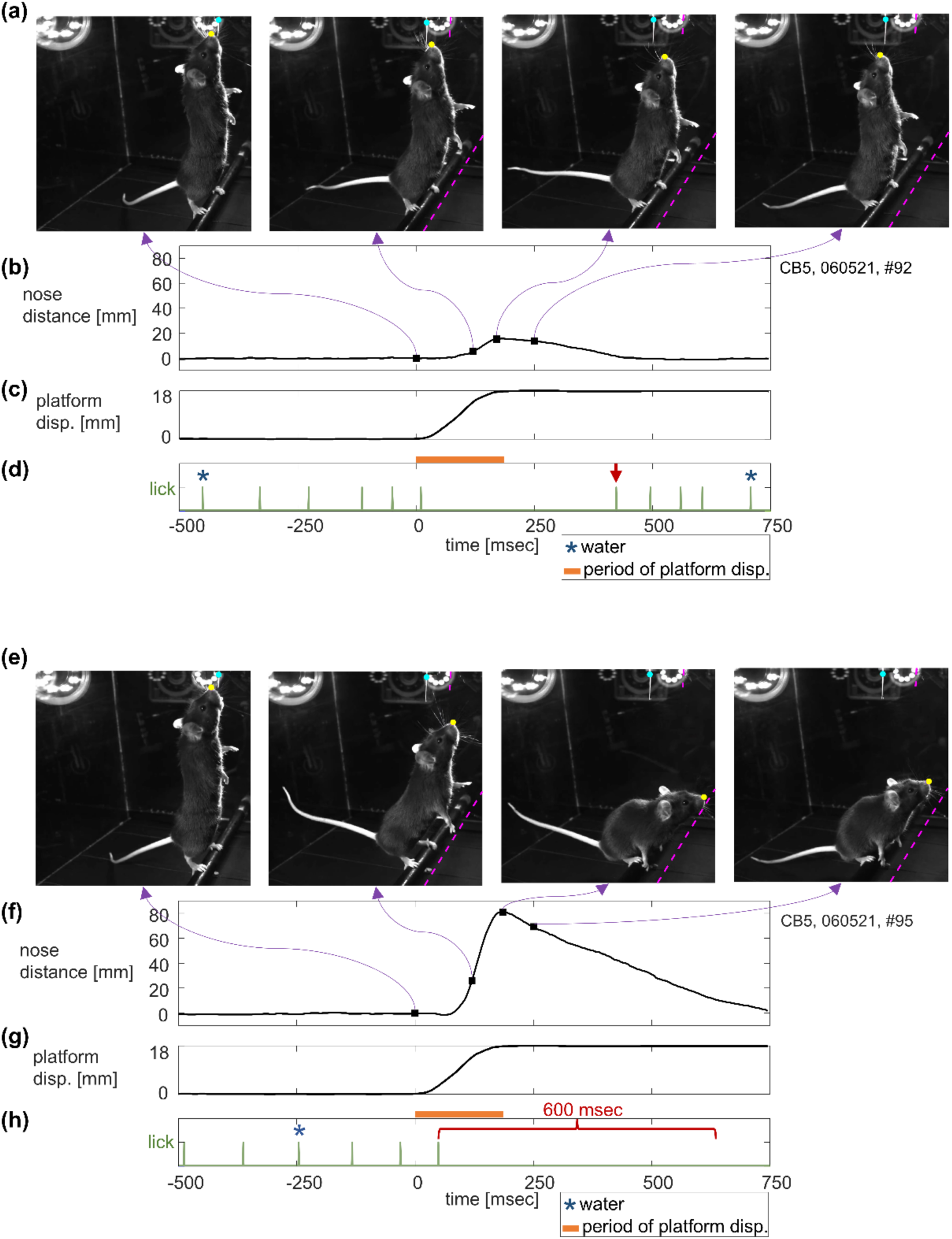
Postural responses to the backward perturbation. This figure shows data from two trials in mouse CB5. (a) Representative snapshots from a recorded video of a successful trial: 1. perturbation onset, 2. 125 msec after perturbation onset, 3. time of maximum nose distance, 4. 250 msec after perturbation onset. Cyan and yellow circles indicate the spout and nose positions that DeepLabCut tracked. Dashed magenta lines indicate the initial positions of the spout and the perch. (b)-(d) Nose distance trace over time from the same trial along with the platform displacement and lick (green) and water (blue) events. Square dots and purple arrows indicating the timings of the snapshots are superimposed on the nose distance trace. The red arrow marks the timing of lick after the perturbation. (e)-(h) Representative snapshots and time courses from a failed trial, depicted in the same way as (a)-(d). The red bracket highlights a 600-msec interval representing the maximum allowable time between consecutive licks, after which the trial was terminated if the mouse failed to lick again.

A trial could terminate in one of two ways: 1) a ‘complete’ trial if the mouse continued licking throughout the full 7.5 seconds; 2) an ‘aborted’ trial if the mouse failed to lick for 600 msec. When the trial was terminated either way, the LED was turned off and the inter-trial interval started.

During the inter-trial interval, the spout did not dispense water even if licking was detected. On each trial, the duration of the inter-trial interval was drawn from a uniform distribution of 10-15 seconds. For aborted trials, 20 seconds was added to the interval duration (additional penalty interval). This lengthy “time out” for aborted trials was enforced to encourage the mouse to get as much water as possible on each trial, rather than to bail early and move on to the next trial. Before a new trial could be started, the mouse had to refrain from licking for 5 seconds. This “no lick” period was included so that the mouse did not lick continuously throughout the inter-trial interval. Additionally, a trial was manually aborted by the experimenter if the mouse faced the wrong direction to make the posture of the mouse more consistent across trials. This manual intervention was effective and standing in the opposite direction rarely occurred during the final few sessions of the pre-training and the full task sessions.

The session ended when the experimenter observed a decline in task engagement, which was indicated by mice reducing or ceasing to initiate trials. Once this occurred, the task was terminated for that day.

All animals underwent a pre-training phase of 13 to 16 sessions, during which task parameters, such as the water timer, were gradually adjusted to their final values, as shown in Figure 2c. One session was conducted per day. Pre-training sessions were either carried out consecutively or with breaks of one to two days between sessions. Mice were considered fully pre-trained after completing 5–6 sessions with the final parameters, demonstrating consistent performance aligned with task requirements. For CB5 and CB6, the pre-training phase was interrupted for two months due to the experimenter’s illness. However, the final 5–6 sessions for these animals were conducted consecutively, without any breaks, immediately before starting the full task.

##### Full task

On top of the trial interval structure that mice learned during pre-training, the sound cue and platform movement were added to the task. The structure of the full task was explained in the subsection *Predictive postural control task* in the Methods section.

#### Perturbation profiles

Though the effect of different perturbation amplitudes was not systematically studied, we explored a few different ones in our pilot experiments to see what was reasonable for the mouse. In what follows, we chose a perturbation size of 18 mm (large) as one that was big enough to be challenging for the mice but not so large as to be impossible to compensate. We also used the perturbation sizes of 7 mm (small, only used for mouse CB10) and 12 mm (medium) as milder ones. The perturbation profiles were determined through pilot experiments (not recorded) for each mouse, which allowed us to adjust parameters from initial estimates based on human studies. These pilot tests provided an empirical foundation for selecting motor settings that would reliably elicit balance responses. For the three platform displacement amplitudes, peak velocity was 60 mm/sec (small, only used for mouse CB10), 100 mm/sec (medium), and 140 mm/sec (large) respectively. Only one amplitude was used for any single session.

### Data Acquisition

#### Video recording

Four video cameras (OptiTrack; NaturalPoint, Corvallis, OR; Table 2) were mounted to surround the behavioral arena (Figure 2b). Mice were videotaped at 200 fps with 1280 x 1024 pixels resolution, and data was saved using OptiTrack recording software, Motive (NaturalPoint). For the present analysis, we used only the data obtained by a single camera.

#### Event recording

The timing of critical trial events, such as LED onset, licks, rewards, and video frames, were marked by digital voltage signals (transistor–transistor logic or TTL) and recorded to the PC via a data acquisition board (USB-6002, National Instruments) controlled by custom scripts written with LabVIEW software (National Instruments). To synchronize the video data with trial events, an eSync 2 device (NaturalPoint) generated a voltage signal at the start of each video frame, and these signals were also recorded by the PC via the data acquisition board.

### Data Processing and Analysis

#### Video tracking using DeepLabCut

A deep learning-based pose estimation system, DeepLabCut (Lauer et al., 2022; Mathis et al., 2018; Nath et al., 2019) version 2.3, was used to track key points of the mouse and the apparatus. From the perspective of motor control, one can hypothesize that the goal of the mouse is to control the location of their tongue close to the end of the lick spout. Thus, one good measure of postural response would be the distance of the control point (tongue) from the target (end of lick spout). Since the tongue was not constantly visible in the videos, we tracked the position of the nose. For the same reason, a visibly distinct part of the spout was tracked instead of the end of the lick spout (e.g., Figure 3a and 3e). Distance between these two tracked points (hereafter, “nose distance”) was used as an index of postural response.

In order to train the DLC network, frames were extracted for manual labeling using the k-means clustering algorithm and manual selection, to reflect the diversity of images. The resulting training dataset consisted of 288 frames (from 36 videos) from all three animals. These manually labeled images were then used to train the weights of a standard pre-trained network (ResNet-50) for 1,030,000 training iterations. 90% of the labeled frames were used to train the DLC network, and performance was validated using the remaining 10%.

After running DeepLabCut on each video file, the output files (CSV file with x/y coordinates of selected features and their corresponding likelihood values) were processed. The nose distance was obtained as the Euclidian distance between the nose and the spout in each video frame. The baseline nose distance was defined by taking the mean of the nose distance of 250 – 2,250 msec before the perturbation onset and was used for subtraction. Any nose distance value exceeding 450 pixels (= 81 mm) was capped at 450 pixels. At this distance, the mice came down from bipedal standing and their front paws were near the height of the perch. Nose distance values larger than this were often contaminated by the motions of the animals that were not the target of interest (e.g., they stepped down from the perch or stepped to the right or left on the perch).

We removed certain trials from the analysis where the nose distance at the perturbation onset was more than 225 pixels (= 40.5 mm) (0.54% of all analyzed trials). A nose distance of 225 pixels or more at the perturbation onset indicates that the mouse was ducking down at the onset of the perturbation. This itself is an interesting behavior as it could indicate that mice are predicting the timing of the perturbation. However, if the initial posture was not standing upright, it did not make sense to compare responses to the perturbation. Therefore, those trials were excluded from the analysis.

We also note that 8 trials (0.36% of all analyzed trials) had to be excluded from the maximum nose distance analysis (explained in the following subsection) because no videos were recorded due to camera failures. The trial events data for these trials were intact and used for the trial outcome analysis.

#### Quantification of postural responses

##### Maximum nose distance

To quantify the postural performance for each trial, we used maximum distance of the nose position from the lick spout during the time window from the onset of the perturbation to 250 msec after the perturbation onset. We looked at other measures such as mean, median, and area under the trace, and found that the main results did not change.

##### Trial outcome

From the reward learning perspective, one can hypothesize that the goal of the mouse is to maximize the amount of water reward that they obtain on a given trial. Thus, we defined another performance index based on rewards that the mouse obtained. We consider a trial to be successful if the animal maintains its posture and continues licking throughout the perturbation. Success is operationally defined by whether the mouse is able to obtain a water droplet from the lick spout after the perturbation (600 msec time window from the last lick). In a given trial, if mice obtained at least one water droplet after the perturbation, we classified the trial as a “success.” In a given trial, if mice obtained no water droplet after the perturbation, we classified it as a “failure”. We call whether a trial was a success or failure the “trial outcome.” Note that if mice failed to lick within the 600-msec interval, the trial was aborted (see Methods; *Predictive postural control task*). Therefore, if mice could not recover to the lick spout quickly enough and lick again after the perturbation, they could no longer obtain any water reward (see Figure 3 for a graphical explanation).

In a successful trial, mice tended to get the maximum number of water droplets (7 droplets per trial) unless they stopped licking during the post-perturbation period (Figure 4). In a failed trial, mice received a few water droplets before perturbation and no reward after the perturbation (Figure 4).

**Figure 4:**
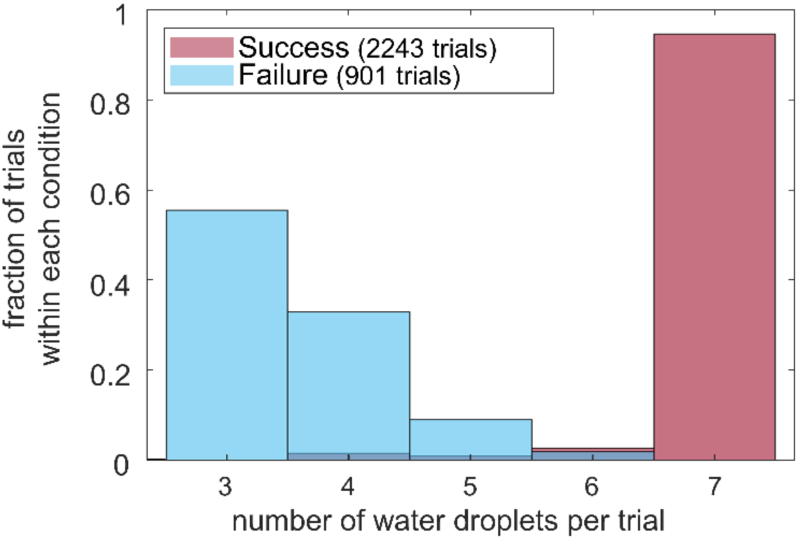
Distribution of number of received water droplets per trial. Distribution of the number of received water droplets per trial for success and failure from all three animals.

Consequently, the total number of water droplets mice received in a failed trial varied depending on the timing of the perturbation.

#### Regression models

We used regression models to analyze the effect of the sound cue and learning on the postural responses.

##### Exponential decay model to predict the maximum nose distance

For each animal, we performed nonlinear regression (using the ‘fitnlm’ function in MATLAB; MathWorks, Natick, MA) to individual trial data, to predict the maximum nose distance based on two input variables - namely, whether a trial was cued or not (indicator variable) and the session index (order of experimental sessions across days). We used an exponential decay model as it is one of the most traditional models of learning and is also used in human postural perturbation studies(Kolb et al., 2004, 2002, 2000) (series of work from Kolb et al.). We did not incorporate the trial index as a variable in this model because we did not see a consistent trend based on it across sessions. We will discuss the effect of it in a separate section.

Equation 1 describes standard decay function:

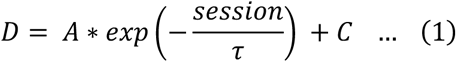

where *D* corresponds to the maximum nose distance and “*session*” corresponds to the session index. *A*, *τ*, and *C* are the coefficients to be fit by the regression, where *A* represents the amplitudes of the decay component, *τ* is the decay time constant across sessions, and *C* is an offset.

To incorporate the effect of the cue, we used the “learning rate model,” in which the cue affects the learning rate (time constant, *τ*, in Equation 1). The learning rate model would indicate that the cue somehow speeds up or slows down the associative process of learning over sessions.

This model can be formulated in the following equation.

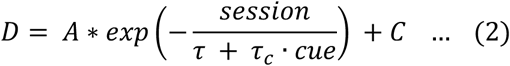

In this equation, “*cue*” is equal to 0 on NOCUE trials and 1 on CUE trials. *A*, *τ*, *τ_c_*, and *C* are the coefficients to be fit by the regression; *τ_c_* represents the effect of the cue on the time constant. A negative value for *τ_c_* would mean that mice learn faster over cued trials, and vice versa.

##### Logistic regression model to predict the trial outcome

For each animal, we performed logistic regression (using the generalized linear model; ‘fitglm’ in MATLAB) to individual trial data, to predict the trial outcome based on whether a trial was cued or not (indicator variable), the session index, and the trial index. Equation 3 describes this relationship:

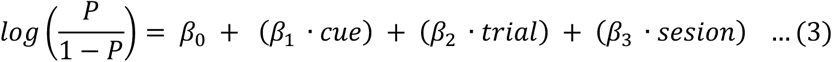

or equivalently,

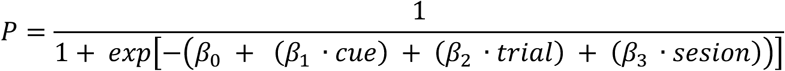

where “*P*” corresponds to the probability of success and *β_0_*, *β_1_*, *β_2_*, and *β_3_* are the coefficients to be fit by the regression. *β_0_* is a coefficient that represents the log-odds of a successful trial without any prior experience of the task or the cue. *β_1_* is a coefficient representing the effect of cue on the trial outcome. *β_1_* is multiplied by “*cue*,” which is equal to 0 on NOCUE trials and 1 on CUE trials. *β_2_* models the effect of trial index on the trial outcome; “*trial*” is the trial index within a session. *β_3_* is the coefficient corresponding to the effect of session index on the trial outcome.

#### Statistical analysis

Data analysis was performed using custom scripts written in MATLAB R2020a (MathWorks). For each animal, one regression model and one logistic regression model were fit using MATLAB’s fitnlm() function and fitglm() function, respectively: one to predict nose distance and one to predict the outcome of the trial.

## Results

The movements of the mice were simultaneously recorded by four cameras (C1 – C4) which were mounted to surround the behavioral arena (Figure 5). In the present work, we analyzed the videos from one camera (C4) as data from a single camera was sufficient to demonstrate the validity of the task. This approach was reliable for our specific postural assessment, given that the animals’ heads remained in a fixed plane relative to the lick spout. In this report, we focus on our analyses using the data collected from camera C4 because it provided the optimal view for our purpose as it was positioned closest to the sagittal plane, offering the clearest perspective for assessing postural changes. The view angle of each camera remained constant with respect to the behavioral apparatus throughout the experiment. The behavioral and task events such as licking, cue onset, and water deliveries were also recorded and synchronized with the video recordings, and this allowed event-based analyses (Figure 2; see Methods for the details).

**Figure 5:**
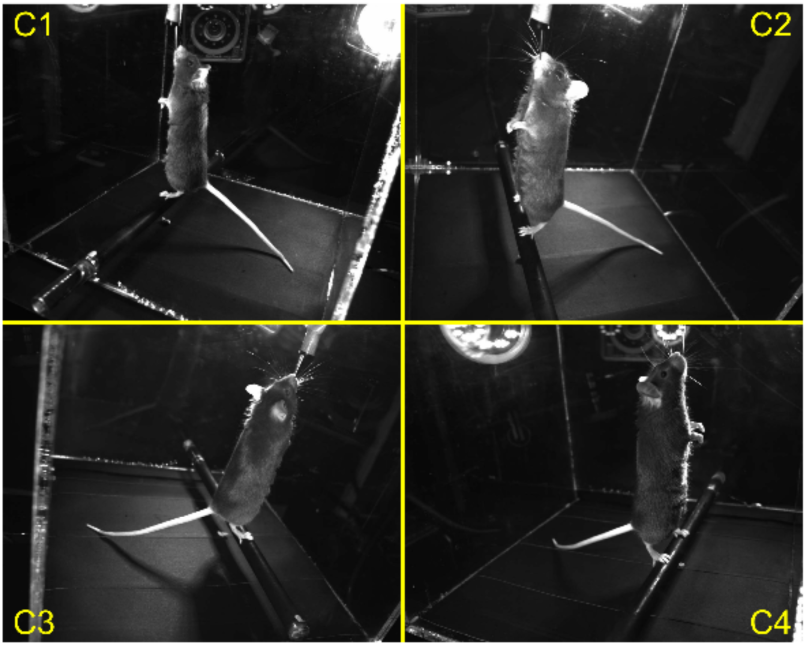
Views from four cameras. An example simultaneous view from four cameras during the task. The analysis in this paper is using C4 camera only.

Here, we will show two quantifications; one using the tracked video data and the other using recorded behavioral and task events, as an example of what one can readily measure using our task. Then, we will demonstrate the measurable effect of cue and learning using these two quantifications. In order to quantify the postural performance for each trial, we used two indices: (1) maximum distance of the nose position from the lick spout during the time window from the onset of the perturbation to 250 msec after perturbation onset (the maximum nose distance) (Figure 6a magenta lines showing the time window), and (2) whether the animal obtained at least one water reward after the end of perturbation (trial outcome).

**Figure 6:**
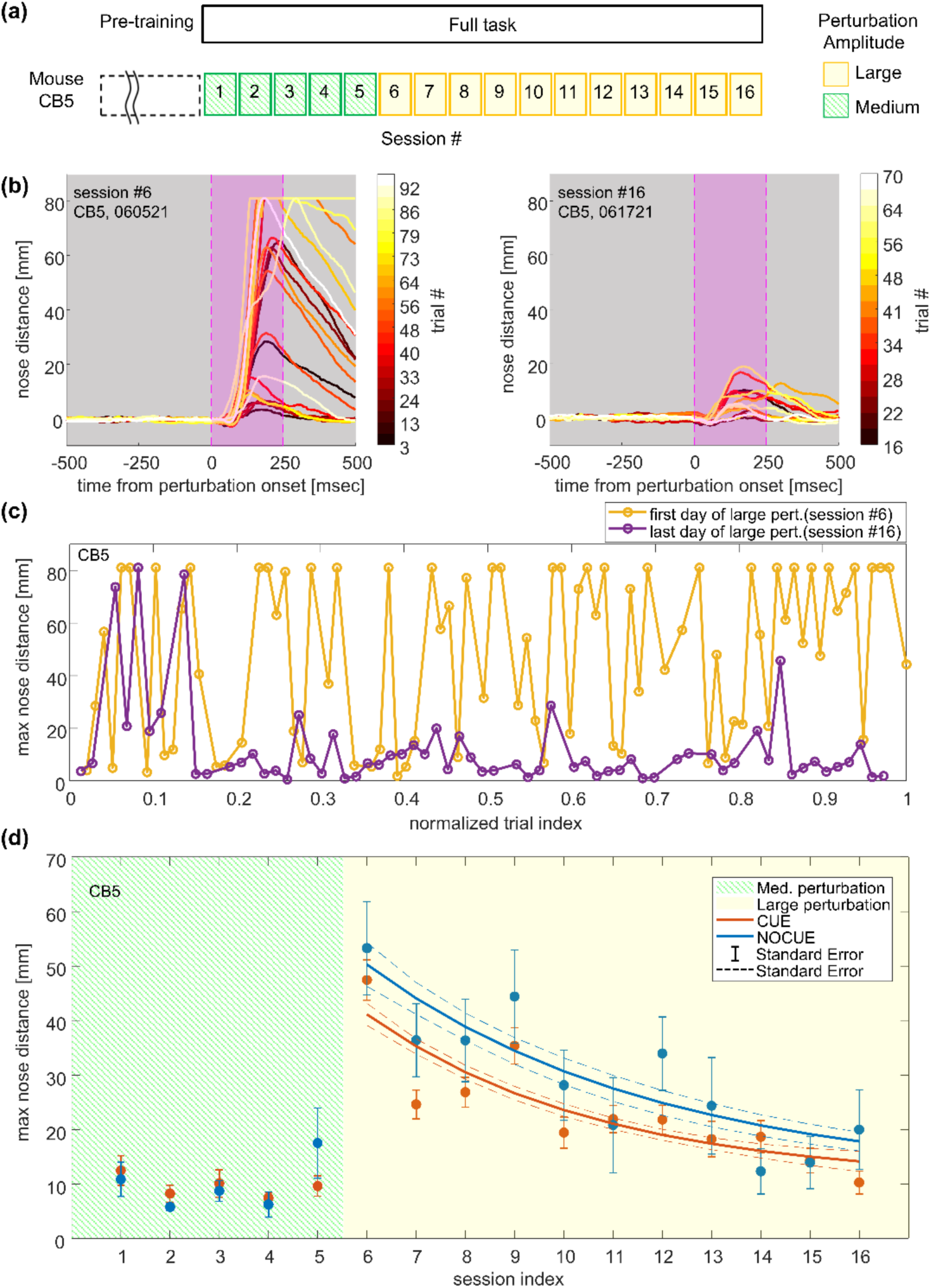
Data analysis based on nose distance. This figure shows data from an example mouse (mouse CB5). (a) Time course of sessions. The platform was moved in one of three amplitudes: 7 (small), 12 (medium), or 18 (large) mm. Only one amplitude was used for an individual session. (b) Nose distance traces for the first session (left) and last session (right) of large (18 mm) disturbance days. Every 3 trials are plotted, and traces are color-coded by trial index. Note that trials 1-12 of the last session were excluded from this figure due to the low quality of DeepLabCut tracking. However, the max nose distances of those trials were manually extracted using ImageJ and used in the following figures and analysis (see Methods). (c) Max nose distances across all trials for the first session (yellow) and last session (purple) of large disturbance days. Trial indexes are normalized by the total number of trials (82 and 63 trials for the first and last session respectively) of the session. (d) Mean and standard errors of the max nose distance across all the sessions. Lines show fit from an exponential decay model (“learning rate model”). Dashed lines represent standard errors for fits.

For illustration purposes, four representative snapshots from a recorded video of a successful trial and a failed trial from one of the cameras are shown in Figure 3a and 3e respectively: Frame 1, perturbation onset; Frame 2, 125 msec after perturbation onset (the middle time point of the time window for calculating maximum nose distance); Frame 3, time of maximum nose distance; Frame 4, 250 msec after perturbation onset (the end of the time window for calculating maximum nose distance). In most of the trials, the backward platform movement produced forward movement of the mouse’s body, which is consistent with the human postural responses to backward perturbations (Horak and Nashner, 1986). Typically, the tail tip and the heels went up with forward movement of the whole body and then went back down as mice recovered to the upright position. See Video 1 (successful trial) and Video 2 (failed trial) for the same representative trials (at 0.25x speed).

### Nose distance

The time series of nose distance of the same representative trials are shown in Figure 3b and 3f with square dots indicating the timing of snapshots along with the time course of platform displacement (Figure 3c and 3g). From these representative traces, we can see that there was about 50 – 100 msec of delay between the onset of the perturbation and the onset of the nose distance displacement. This kind of delay is consistent with what was observed in the human postural paradigm (Welch and Ting, 2014). The representative traces also showed that the nose distance peaked near the end of the perturbation and gradually recovered to the baseline as mice reassumed an upright posture. The descending recovery slopes were more gradual than the ascending slopes. We also note that, although not depicted in these example trial traces, there were cases where we observed changes in nose distance and other kinematics before the perturbation onset. This could be a sign of anticipation or learning and could also modulate the postural responses to the perturbation.

Figure 6 shows the nose distance data from this mouse. The mouse experienced medium-amplitude perturbations on session #1 through #5, and large-amplitude perturbations on session #6 through #16 (Figure 6a). In Figure 6b, multiple nose distance traces for the first (left) and last (right) sessions of the large perturbation days are shown. The traces are aligned to the perturbation onset, and every 3rd trial is plotted for visualization purposes. We observed some variability in the onset of nose displacement and the timing of the peak, but the overall time course is consistent: peaking near the end of the perturbation and more gradual recovery.

The maximum nose distance fluctuated across trials (Figure 6b, c). However, on average, the last session had smaller maximum nose distances compared to the first session of the large perturbation days. In one of the example sessions shown in Figure 6c, the maximum nose distance was relatively large at the beginning of the session but was reduced throughout the rest of the session (session #16, the last day of large perturbation). On the other hand, in the other session shown in Figure 6c, a trend across trials was barely apparent (session #6, the first day of large perturbation). The overall postural performance for each session was summarized by the mean and standard error of maximum nose distances (Figure 6d for mouse CB5).

### Trial outcome

The timing of licks and water deliveries of the same representative trials are shown in Figure 3d and 3h (mouse CB5). The mouse licked relatively constantly during the pre-perturbation period, and then could not lick for a certain amount of time due to the effect of postural perturbation.

Figure 7 shows the trial outcome data from this mouse (mouse CB5). The sliding window proportion of success and every trial outcome in the first and last session of the large perturbation sessions are shown in Figure 7a top and bottom respectively. Overall, the last session had a higher success proportion than the first session. The overall postural performance of each session was summarized by the proportion of successful trials over all the trials (Figure 7b), and error bars represent 95% confidence intervals estimated based on the binomial distribution.

**Figure 7:**
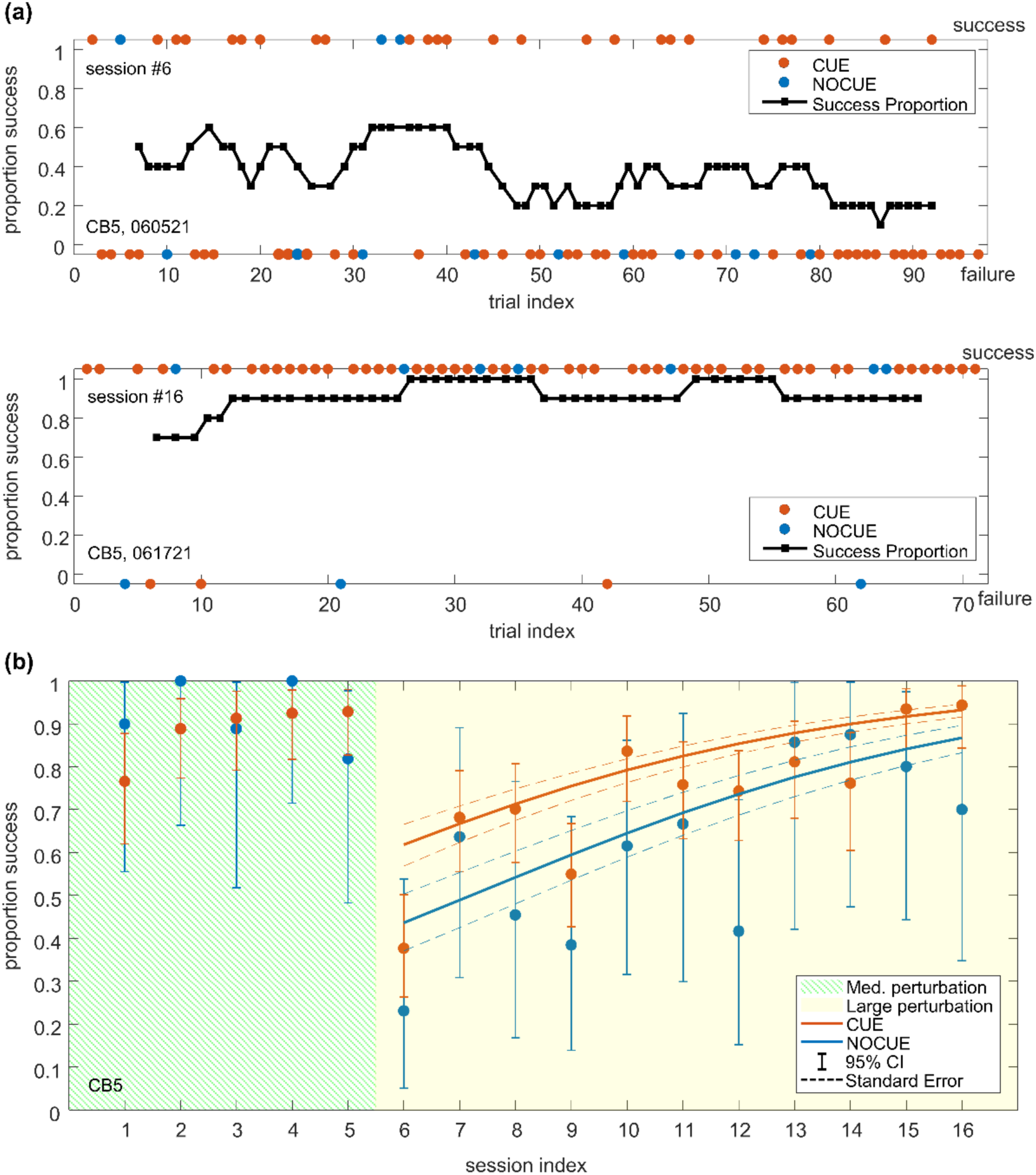
Data analysis based on success proportion. This figure shows data from an example mouse (mouse CB5). (a) The sliding window proportion of successes (sliding window size = 10, black) and every trial outcome (colored circles) in the first (top) and last (bottom) session of the large perturbation sessions. (b) The success proportion with 95% confidence intervals across all sessions. Solid lines show fits from logistic regression for CUE and NOCUE conditions. Dashed lines represent standard errors for fits.

### Measurable effect of cue and learning

To analyze the effect of the sound cue and learning on the postural responses, we used regression models (see Methods). These models allowed us to assess the strength of the relationship between postural responses and several predictor variables. This allowed us to simultaneously test whether animals’ performance improved both within and between sessions and to determine whether the presence of a predictive cue affected performance while controlling for the other variables.

For this example mouse (mouse CB5), the exponential function (the predictor variables were: whether a trial was cued or not [indicator variable] and the session index; see Methods for details) described the trend over sessions in the maximum nose distance relatively well (adjusted *R^2^*= 0.13; Figure 6d, *n* = 795 trials, showing the fitted lines for the learning rate model; see Methods for the details of the model). This suggests that learning across sessions can be well characterized by an exponential decay function for maximum nose distance in this mouse. The model had a negative value for the cue-related coefficient (*τ_c_* = −1.21, p = 0.058 in the learning rate model), which suggests that the cue works in the direction of *improving* the postural response. Table 3 shows the coefficients and p-values of the model for this mouse.

**Table 3:** Exponential Decay Regression Models Coefficients and Statistics.

**Mouse CB5 (learning rate model, AIC =7270.6)**
|  | **Coefficients** | **Standard Errors** | **P-Values** |
| --- | --- | --- | --- |
| A | 110 | 35 | 0.0016* |
| $\tau$ | 5.9 | 2.0 | 0.0036* |
| $\tau_c$ | -1.2 | 0.6 | 0.058 |
| C | 10.5 | 4.6 | 0.023* |

\* indicates significant value (p <0.05)

For the same mouse (CB5), we also examined the effect of within-session and across-session learning, and the effect of cue on the trial outcome using logistic regression (the predictor variables were: whether a trial was cued or not [indicator variable] and the session index, and the trial index within a session; see Methods for details), and found that the cue and the session index had significant positive regression coefficients (*e^β1^*= 2.10, p = 0.00073; *e^β3^* = 1.24, p <0.0001; Table 4) and the trial index had a negative regression coefficient (*e^β2^* = 0.99, p = 0.033; Table 4). Exponentiated coefficients represent the effect size of the cue in terms of the odds-ratio in logistic regression. Thus, these results indicate that the odds of a successful trial are 2.10-fold higher with the cue compared to without the cue and that one increment in the session index multiplies the odds of a successful trial by 1.24 and one increment in the trial index multiplies the odds of a successful trial by 0.99 for this mouse (Figure 7b, *n* = 797 trials; see also Table 4).

**Table 4:**
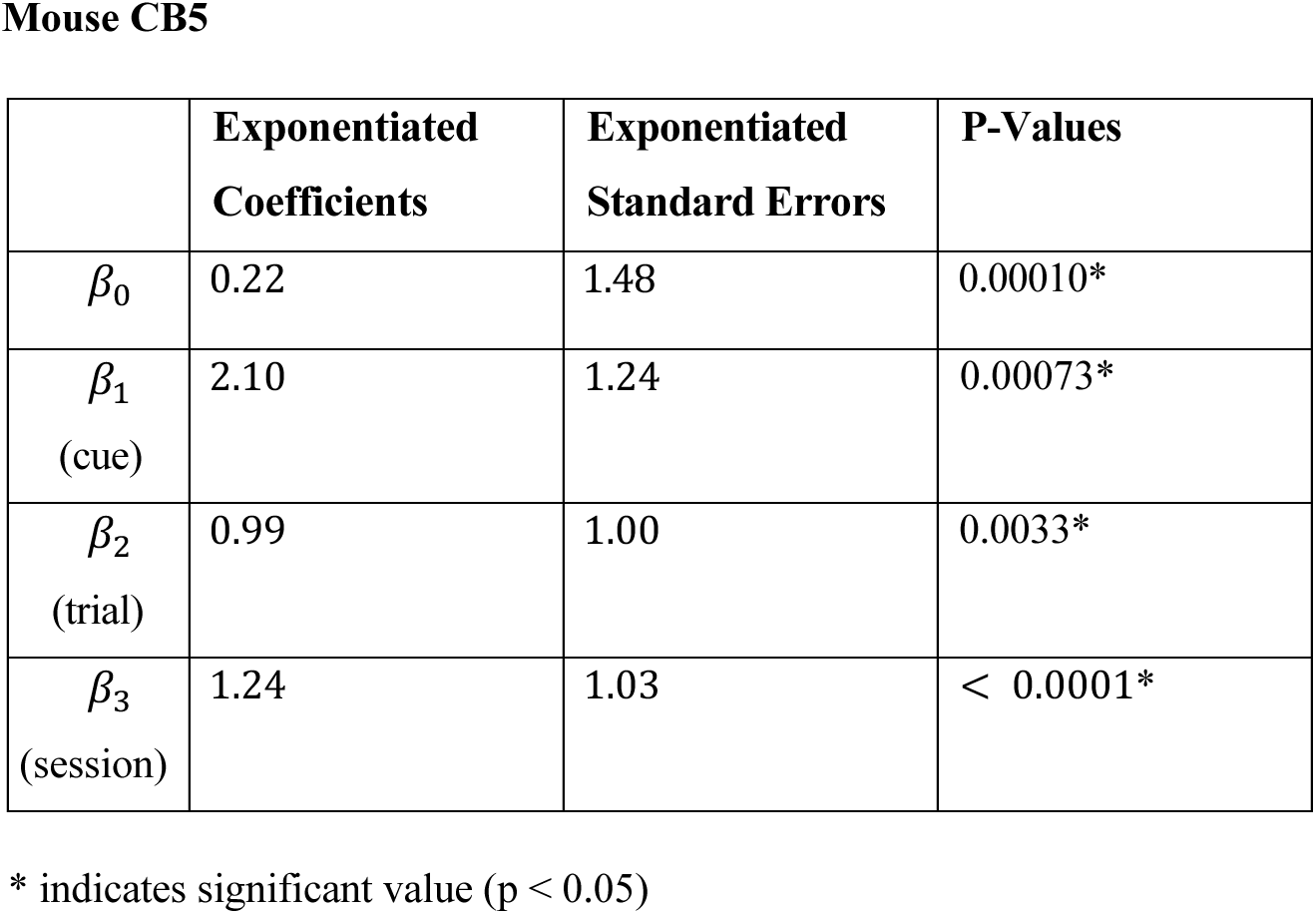
Logistic Regression Model Coefficients and Statistics.

**Mouse CB5**
|  | **Exponentiated**<br>**Coefficients** | **Exponentiated**<br>**Standard Errors** | **P-Values** |
| --- | --- | --- | --- |
| $\beta_0$ | 0.22 | 1.48 | 0.00010* |
| $\beta_1$<br>(cue) | 2.10 | 1.24 | 0.00073* |
| $\beta_2$<br>(trial) | 0.99 | 1.00 | 0.0033* |
| $\beta_3$<br>(session) | 1.24 | 1.03 | < 0.0001* |

\* indicates significant value (p < 0.05)

## Discussion

In this study, we established a postural control experimental paradigm for mice that enables studies of mechanisms underlying predictive postural control. Using a small sample of animals, we show that a combination of video recordings and event data can be used to study postural responses to perturbations between predictable and unpredictable conditions. We utilized an open-source microcontroller (Arduino) for their versatile I/O capabilities and adaptability, allowing seamless integration with neural recording/manipulation devices and easy modification for various postural research tasks in the future.

There are a few limitations and potential improvements to be considered. First, as in any task that uses operant conditioning, the number of trials per session (approximately 100 trials across 3 different trial types) was limited by satiety, which reduces the animals’ motivation to continue performing the task. In our experiments, the total water volume consumed during a session was comparable to that in previous studies (Andermann et al., 2010; Histed et al., 2012; Kim et al., 2020). However, we think there is considerable room to reduce the total number of water rewards per trial, and thus increase the total number of trials within a session. Even larger gains might be achieved by using a non-satiating reward, such as intracranial self-stimulation (Carlezon and Chartoff, 2007; Olds and Milner, 1954). Second, while we chose 75% CUE trials to strengthen cue-movement associations, the limited number of NOCUE trials reduced our ability to detect within-session effects of the cue. Future studies should more closely balance the proportion of CUE and NOCUE trial types for both learning and statistical power. Finally, we observed decreased engagement in later trials through longer initiation times (Figure 8), a common issue in restriction-based behavioral experiments (Berridge, 2004; Ortiz et al., 2020). This could be addressed by implementing online monitoring of trial initiation times, introducing rest periods, or adapting the task for home-cage use (e.g. (Poddar et al., 2013)). Additionally, incorporating cue-only trials (where the auditory cue is presented without perturbation) would be important to further investigate the biological process of how the cue induces the beneficial effect on postural responses - if the cue alone elicits postural responses, this would strongly support associative learning (Campbell et al., 2009).

**Figure 8:**
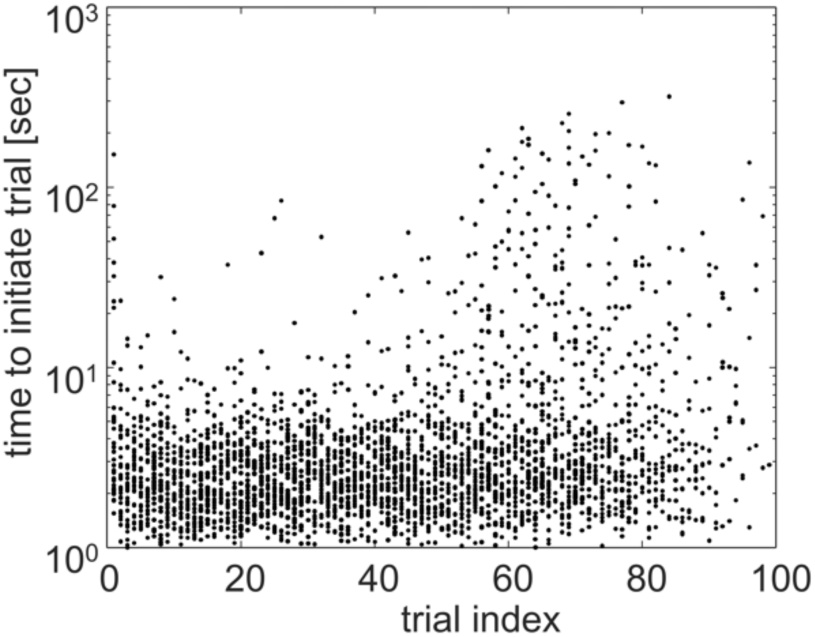
Time to initiate trial. Time to initiate the trial for each trial from sessions in which this information was available (all sessions for CB5 and CB6, session #1-5 for CB10) from all three mice are superimposed. Note that each session has a different number of trials. Trials in which the time to initiate is shorter than 1 sec are rare and omitted from this plot.

While mice offer powerful neural recording and manipulation capabilities, important species differences must be considered when comparing to human balance studies. The mouse tail significantly influences balance through multiple mechanisms: adjusting whole-body center of mass, generating compensatory torque, and when floor-touching, providing mechanical support, wider base of support, and sensory feedback. Our observations showed that mice frequently touch the floor with their tails during quiet bipedal standing and make dynamic tail movements during perturbations, suggesting future studies should consider tail tethering and force measurement to isolate other body parts’ contributions to balance (Funato et al., 2017). Additionally, differences in segment alignment and body mass proportions between mice and humans highlight the need for caution in comparative research. Rats in bipedal stance exhibit flexed segment alignment compared to humans’ nearly aligned posture (Funato et al., 2017), and their trunk bending during postural responses to floor rotation differs from humans possibly due to proportional differences in trunk mass and length (Konosu et al., 2021). Although our study did not analyze segment kinematics, similar traits are expected in mice. Despite these biomechanical differences, the inherent instability of mouse bipedal standing necessitates neural control, making them valuable models for studying fundamental postural control mechanisms.

Various paradigms have been developed to study postural control in rodents, ranging from general balance assessments to more sophisticated models. Traditional methods such as the righting reflex test (Pellis, 1996; Whishaw and Kolb, 2020; Yarnell et al., 2016), rotarod test (Crawley, 2007; Dunham and Miya, 1957), and balance beam test (Carter et al., 2001; Luong et al., 2011) provide insights into overall balance but lack specificity on postural control mechanisms. Other relevant studies include Murray et al., 2018 who combined EMG recordings and optogenetics to show distinct neural populations in the lateral vestibular nucleus governing balance during beam traversal with lateral perturbations, and Lee et al., 2015 who demonstrated cerebellar adaptation in rats during a suspended dowel task, revealing how Purkinje cells encode head angle and adapt to filter out self-generated sensory inputs. These studies provide unique opportunities to investigate neural circuits and molecular mechanisms underlying balance control, complementing human research and offering insights that may be difficult to obtain from human subjects alone. Recently, Konosu et al., 2024, 2021 established a paradigm for bipedally standing rats subjected to floor-tilting perturbations preceded by visual cues, showing diminished postural responses with repeated perturbations, indicating plasticity in postural control. Building on this, our study investigated the role of prediction by comparing responses to backward floor translations that were either unpredictable or preceded by auditory cues. Several differences distinguish our work from this previous research. While their study employed a flat-floor setup, we used a perch-balancing task. The use of perch allowed us to constrain the locations of the animal relative to the water spout, thus to reduce behavioral variability. We note, however, that this may engage different muscle groups due to the demands of grasping. Additionally, the time scales differed; the earlier study conducted sequential cued trials within a single day, whereas our research extended over two weeks, incorporating both cued and unpredictable perturbations. This design allowed us to directly measure the effects of the auditory cue on learning and postural responses both within and across days. Furthermore, the types of perturbations varied: the earlier study used rotational perturbations, which simplify kinematic analysis but involve more complex mechanical setups, whereas we employed translational perturbations, which simplify the mechanical design but require accounting for horizontal movement during analysis.

Future developments of this study could explore posture control dynamics through deeper analysis of kinematics and kinetics, combined with neural circuit investigation. Specific advancements could include center of mass analysis which is often used in human postural studies (Peterka, 2002; Ting, 2007; Van Wouwe et al., 2021; Welch and Ting, 2014), three-dimensional kinematics from multiple cameras (Dunn et al., 2021; Karashchuk et al., 2021; Marshall et al., 2020; Nath et al., 2019), and unsupervised movement clustering (Marshall et al., 2020; Wiltschko et al., 2020, 2015) to reveal novel postural strategies. Additionally, incorporating EMG recordings and force measurements would help detect subtle postural adjustments such as muscle co-contraction and toe gripping that are invisible in kinematic data alone.

To conclude, we have established a mouse experimental paradigm to study predictive postural control, an understudied aspect of motor control. By combining modern computer vision and machine learning capabilities with advanced neuroscience tools available in mice, this paradigm opens the door to understanding the neural circuits underlying predictive postural control.

## Acknowledgments

We thank Akira Konosu and all Uchida and Yanagihara lab members for their discussion and feedback. We also would like to acknowledge the support of Ed Soucy, Brett Graham, and the Center for Brain Science Neurotechnology Core for their help in building the experimental apparatus, and Kohei Yamaji for his help in illustrations. This work was supported by Simons Collaboration of the Global Brain (N.U.), Grants-in-Aid for for Scientific Research (C) (18K10955) funded by the Ministry of Education, Culture, Sports, Science, and Technology of Japan (D.Y.), gift from the Ilfield Fund to (R.T.B.), Murata Overseas Scholarship (Y.D.), and Masason Foundation (Y.D.).

## References

Andermann ML, Kerlin AM, Reid RC (2010) Chronic cellular imaging of mouse visual cortex during operant behavior and passive viewing. Front Cell Neurosci 4:3.

Bagnall MW, McLean DL (2014) Modular organization of axial microcircuits in zebrafish. Science 343:197–200.

Bastian AJ (2006) Learning to predict the future: the cerebellum adapts feedforward movement control. Curr Opin Neurobiol 16:645–649.

Berridge KC (2004) Motivation concepts in behavioral neuroscience. Physiol Behav 81:179–209.

Campbell AD, Dakin CJ, Carpenter MG (2009) Postural responses explored through classical conditioning. Neuroscience 164:986–997.

Carlezon WA Jr, Chartoff EH (2007) Intracranial self-stimulation (ICSS) in rodents to study the neurobiology of motivation. Nat Protoc 2:2987–2995.

Carter RJ, Morton J, Dunnett SB (2001) Motor coordination and balance in rodents. Curr Protoc Neurosci Chapter 8:Unit 8.12.

Coelho DB, Luis, Teixeira A (2017) Cognition and balance control: does processing of explicit contextual cues of impending perturbations modulate automatic postural responses? Exp Brain Res 235:2375–2390.

Cohen JY, Haesler S, Vong L, Lowell BB, Uchida N (2012) Neuron-type-specific signals for reward and punishment in the ventral tegmental area. Nature 482:85–88.

Crawley JN (2007) What’s wrong with my mouse?: Behavioral phenotyping of transgenic and knockout mice. Hoboken, NJ, USA: Wiley.

Dakin CJ, Bolton DAE (2018) Forecast or Fall: Prediction’s Importance to Postural Control. Front Neurol 9:924.

Deliagina TG, Orlovsky GN (2002) Comparative neurobiology of postural control. Curr Opin Neurobiol 12:652–657.

Dunham NW, Miya TS (1957) A note on a simple apparatus for detecting neurological deficit in rats and mice. J Am Pharm Assoc Am Pharm Assoc 46:208–209.

Dunn TW, Marshall JD, Severson KS, Aldarondo DE, Hildebrand DGC, Chettih SN, Wang WL, Gellis AJ, Carlson DE, Aronov D, Freiwald WA, Wang F, Ölveczky BP (2021) Geometric deep learning enables 3D kinematic profiling across species and environments. Nat Methods 18:564–573.

Eshel N, Tian J, Bukwich M, Uchida N (2016) Dopamine neurons share common response function for reward prediction error. Nat Neurosci 19:479–486.

Fujio K, Obata H, Kawashima N, Nakazawa K (2016) The effects of temporal and spatial predictions on stretch reflexes of ankle flexor and extensor muscles while standing. PLoS One 11.

Funato T, Sato Yota, Fujiki S, Sato Yamato, Aoi S, Tsuchiya K, Yanagihara D (2017) Postural control during quiet bipedal standing in rats. PLoS One 12.

Histed MH, Carvalho LA, Maunsell JHR (2012) Psychophysical measurement of contrast sensitivity in the behaving mouse. J Neurophysiol 107:758–765.

Horak FB, Diener HC (1994) Cerebellar control of postural scaling and central set in stance. J Neurophysiol 72:479–493.

Horak FB, Diener HC, Nashner LM (1989) Influence of central set on human postural responses. J Neurophysiol 62:841–853.

Horak FB, Nashner LM (1986) Central programming of postural movements: adaptation to altered support-surface configurations. J Neurophysiol 55:1369–1381.

Jacobs JV, Fujiwara K, Tomita H, Furune N, Kunita K, Horak FB (2008) Changes in the activity of the cerebral cortex relate to postural response modification when warned of a perturbation. Clin Neurophysiol 119:1431–1442.

Jacobs JV, Horak FB (2007) Cortical control of postural responses. J Neural Transm 114:1339–1348.

Karashchuk P, Rupp KL, Dickinson ES, Walling-Bell S, Sanders E, Azim E, Brunton BW, Tuthill JC (2021) Anipose: A toolkit for robust markerless 3D pose estimation. Cell Rep 36:109730.

Kilkenny C, Browne WJ, Cuthill IC, Emerson M, Altman DG (2010) Improving bioscience research reporting: the ARRIVE guidelines for reporting animal research. PLoS Biol 8:e1000412.

Kim HR, Malik AN, Mikhael JG, Bech P, Tsutsui-Kimura I, Sun F, Zhang Y, Li Y, Watabe-Uchida M, Gershman SJ, Uchida N (2020) A unified framework for dopamine signals across timescales. Cell 183:1600–1616.e25.

Kolb FP, Lachauer S, Maschke M, Timmann D (2004) Classically conditioned postural reflex in cerebellar patients. Exp Brain Res 158:163–179.

Kolb FP, Lachauer S, Maschke M, Timmann D (2002) Classical conditioning of postural reflexes. Pflugers Arch 445:224–237.

Kolb FP, Timmann D, Baier PC, Diener HC (2000) Classically conditioned withdrawal reflex in cerebellar patients. 2. Impaired unconditioned responses. Exp Brain Res 130:471–485.

Konosu A, Funato T, Matsuki Y, Fujita A, Sakai R, Yanagihara D (2021) A Model of Predictive Postural Control Against Floor Tilting in Rats. Front Syst Neurosci 15:785366.

Konosu A, Matsuki Y, Fukuhara K, Funato T, Yanagihara D (2024) Roles of the cerebellar vermis in predictive postural controls against external disturbances. Sci Rep 14:3162.

Lauer J, Zhou M, Ye S, Menegas W, Schneider S, Nath T, Rahman MM, Di Santo V, Soberanes D, Feng G, Murthy VN, Lauder G, Dulac C, Mathis MW, Mathis A (2022) Multi-animal pose estimation, identification and tracking with DeepLabCut. Nat Methods 19:496–504.

Lee RX, Huang J-J, Huang C, Tsai M-L, Yen C-T (2015) Plasticity of cerebellar Purkinje cells in behavioral training of body balance control. Front Syst Neurosci 9:113.

Lockhart DB, Ting LH (2007) Optimal sensorimotor transformations for balance. Nat Neurosci 10:1329–1336.

Luong TN, Carlisle HJ, Southwell A, Patterson PH (2011) Assessment of motor balance and coordination in mice using the balance beam. J Vis Exp.

Macpherson JM, Everaert DG, Stapley PJ, Ting LH (2007) Bilateral vestibular loss in cats leads to active destabilization of balance during pitch and roll rotations of the support surface. J Neurophysiol 97:4357–4367.

Marshall JD, Aldarondo DE, Dunn TW, Wang WL, Berman GJ, Ölveczky BP (2020) Continuous Whole-Body 3D Kinematic Recordings across the Rodent Behavioral Repertoire. Neuron.

Mathis A, Mamidanna P, Cury KM, Abe T, Murthy VN, Mathis MW, Bethge M (2018) DeepLabCut: markerless pose estimation of user-defined body parts with deep learning. Nat Neurosci 21:1281–1289.

McChesney JW, Sveistrup H, Woollacott MH (1996) Influence of auditory precuing on automatic postural responses. Exp Brain Res 108:315–320.

Mochizuki G, Sibley KM, Esposito JG, Camilleri JM, McIlroy WE (2008) Cortical responses associated with the preparation and reaction to full-body perturbations to upright stability. Clin Neurophysiol 119:1626–1637.

Murray AJ, Croce K, Belton T, Akay T, Jessell TM (2018) Balance Control Mediated by Vestibular Circuits Directing Limb Extension or Antagonist Muscle Co-activation. Cell Rep 22:1325–1338.

Nath T, Mathis A, Chen AC, Patel A, Bethge M, Mathis MW (2019) Using DeepLabCut for 3D markerless pose estimation across species and behaviors. Nat Protoc 14:2152–2176.

Olds J, Milner P (1954) Positive reinforcement produced by electrical stimulation of septal area and other regions of rat brain. J Comp Physiol Psychol 47:419–427.

Ortiz AV, Aziz D, Hestrin S (2020) Motivation and Engagement during Visually Guided Behavior. Cell Rep 33:108272.

Pellis SM (1996) Righting and the modular organization of motor programs. Measuring movement and locomotion: From invertebrates to humans 111–130.

Percie du Sert N et al. (2020) Reporting animal research: Explanation and elaboration for the ARRIVE guidelines 2.0. PLoS Biol 18:e3000411.

Peterka RJ (2002) Sensorimotor integration in human postural control. J Neurophysiol 88:1097–1118.

Poddar R, Kawai R, Ölveczky BP (2013) A fully automated high-throughput training system for rodents. PLoS One 8:e83171.

Pruszynski JA, Scott SH (2012) Optimal feedback control and the long-latency stretch response. Exp Brain Res 218:341–359.

Schepens B, Drew T (2004) Independent and convergent signals from the pontomedullary reticular formation contribute to the control of posture and movement during reaching in the cat. J Neurophysiol 92:2217–2238.

Silva MB, Coelho DB, de Lima-Pardini AC, Martinelli AR, Baptista T da S, Ramos RT, Teixeira LA (2015) Precueing time but not direction of postural perturbation induces early muscular activation: Comparison between young and elderly individuals. Neurosci Lett 588:190–195.

Straka H, Baker R (2013) Vestibular blueprint in early vertebrates. Front Neural Circuits 7:182.

Sugioka T, Tanimoto M, Higashijima S-I (2023) Biomechanics and neural circuits for vestibular-induced fine postural control in larval zebrafish. Nat Commun 14:1217.

Ting LH (2007) Dimensional reduction in sensorimotor systems: a framework for understanding muscle coordination of posture. Prog Brain Res 165.

Van Wouwe T, Ting LH, De Groote F (2021) Interactions between initial posture and task-level goal explain experimental variability in postural responses to perturbations of standing balance. J Neurophysiol 125:586–598.

Welch TDJ, Ting LH (2014) Mechanisms of motor adaptation in reactive balance control. PLoS One 9:e96440.

Whishaw IQ, Gorny B, Tran-Nguyen LT, Castañeda E, Miklyaeva EI, Pellis SM (1994) Making two movements at once: impairments of movement, posture, and their integration underlie the adult skilled reaching deficit of neonatally dopamine-depleted rats. Behav Brain Res 61:65–77.

Whishaw IQ, Kolb B (2020) Analysis of behavior in laboratory rats In: The Laboratory Rat, pp215–242. Elsevier.

Wiltschko AB, Johnson MJ, Iurilli G, Peterson RE, Katon JM, Pashkovski SL, Abraira VE, Adams RP, Datta SR (2015) Mapping Sub-Second Structure in Mouse Behavior. Neuron 88:1121–1135.

Wiltschko AB, Tsukahara T, Zeine A, Anyoha R, Gillis WF, Markowitz JE, Peterson RE, Katon J, Johnson MJ, Datta SR (2020) Revealing the structure of pharmacobehavioral space through motion sequencing. Nat Neurosci 23:1433–1443.

Wolpert DM, Flanagan JR (2001) Motor prediction. Curr Biol 11:R729–32.

Yakovenko S, Drew T (2009) A Motor Cortical Contribution to the Anticipatory Postural Adjustments That Precede Reaching in the Cat. J Neurophysiol 102:853–874.

Yarnell AM, Barry ES, Mountney A, Shear D, Tortella F, Grunberg NE (2016) The Revised Neurobehavioral Severity Scale (NSS-R) for rodents. Curr Protoc Neurosci 75:9.52.1–9.52.16.

